# Multiple within species comparisons show Tanganyikan cichlid fish have larger brains in less structurally complex habitats

**DOI:** 10.1101/2024.12.06.627222

**Authors:** Bin Ma, Weiwei Li, Zitan Song, Stefan Fischer, Etienne Lein, Arne Jungwirth, Alex Jordan

## Abstract

Many studies have found a link between higher habitat structural complexity and increased relative brain size in vertebrates. Here we explore this relationship in a multi-species comparison, comparing ten species of wild cichlids that differ in their social and territorial behaviour, but which occur across four ecologically similar but structurally diverse rocky habitats. This design allows us to perform repeated intra-specific comparisons, avoiding confounds associated with comparisons across species boundaries. We sampled 147 fish, analysing brain size and architecture while controlling for body mass and species-specific variability and compared this with habitat complexity, quantified using underwater video and three-dimensional reconstructions. Our results challenge the Clever Foraging Hypothesis (CFH), which posits that greater habitat complexity correlates with larger brain sizes. Contrary to CFH, fish from the least complex habitat had significantly larger brains. Additionally, brain architecture analysis indicated a significant enlargement of the cerebellum in fish from less complex habitats, whereas the hypothalamus showed a non-significant negative trend. Taken together, these results indicate that lower habitat complexity may impose higher cognitive demands on spatial memory and navigation due to limited refuges and increased predation risk. This study highlights the need to reconsider the assumed linear positive relationship between environmental complexity and brain development, suggesting that simpler environments might also impose significant cognitive and ecological challenges that drive brain evolution. Our findings underscore the importance of considering intra-species variability and the specific ecological and cognitive demands of different habitats in studies of brain evolution.

## Introduction

Vertebrate species exhibit substantial variation in brain size, with larger brains often argued to be correlated with enhanced cognitive abilities (Benson-Amram et al., 2016; Deaner et al., 2007; Jerison, 1975; MacLean et al., 2014; Reader et al., 2011). The "Ecological Intelligence Hypothesis (EIH)" (Rosati, 2017; Sol, 2008; Sol et al., 2005) posits that animal cognition has evolved as a response to environmental challenges, particularly in relation to foraging (Barton & Dunbar 1997, Clutton-Brock & Harvey 1980). In support of this hypothesis, many studies have found that differences in brain volume positively correlate with differences in habitat complexity (di Porzio, 2020; Gemma E. White et al., 2015; Safi & Dechmann, 2005; Sobrero et al., 2016). These numerous findings support a dominant theme in comparative neuroanatomy: relatively larger brains can provide the additional cognitive processing power necessary to effectively exploit more complex habitats (Chittka & Niven, 2009; di Porzio, 2020).

A common challenge in many of these studies is the unavoidable conflation of multiple effects. In many cases a comparison between species also entails a difference in ecology, overall habitat type, or life-history. Ideally, a comparison could be conducted on phylogenetically closely related species that occur in a range of habitat complexity in the same general ecotype, minimising confounds. Of even greater value would be a comparison of multiple species that are spatially confined to each habitat, i.e. are not itinerant or grazing species that spend time in multiple different habitats. To this end, fishes emerge as a promising study clade as they possess highly plastic brains in comparison to mammals (K. Kotrschal et al., 1998), facilitating effective analysis at an ecological timescale. As in other vertebrates, studies on fish have frequently demonstrated the positive correlation between environmental complexity and brain size, supporting the idea that environmental complexity demands enhanced neuroanatomical adaptations (DePasquale et al., 2016; K. Kotrschal et al., 1998; P. J. Park & Bell, 2010; Pollen et al., 2007; Shumway, 2008, 2010; White & Brown, 2015). For example, a study of Lake Tanganyika’s Ectodine cichlids found that species living in more complex environments, such as rocky habitats, have larger brain volumes (Pollen et al., 2007). Consequently, a simplified version of EIH, referred to as Clever Foraging Hypothesis (CFH) has been proposed (Newaz I. Ahmed 2016, Park & Bell 2010). CFH predicts that organisms living in more complex environments should have larger brains. (George F. Striedter et al., 2005; P. Park & Bell, 2010).

In this study, we aim to examine the relationship between environmental complexity and brain size by performing multiple intra-species comparisons using wild individuals and direct measurements of habitat at the time of observation, thereby avoiding confounds associated with inter-species differences or use of e.g. aquarium trade collections of uncertain providence. We implemented three key aspects in this study; First, we selected four habitats that are ecologically similar (all rocky habitats) but differ in their structural complexity. We quantified their complexity by 3D reconstructions of the lake bottom; which allowed us to assess complexity not only by rugosity but also substrate homogeneity through topographical features. Second, to minimize the influence of dispersal, we selected ten Tanganyikan cichlid species, seven from the tribe *Lamprologini* that exhibit strong territoriality and philopatry, which provides a stable basis for assessing the effects of habitat structural complexity. In contrast, *Eretmodus cyanosticus*, *Interochromis lookii*, and *Tropheus moorii* are non-territorial and exhibit itinerant behaviours, allowing us to explore whether these species are also sensitive to variations in habitat complexity. Finally, all species occurred in all habitats we sampled, allowing us not only to perform interpopulation comparisons but also to test within each species for generalizability.

We ask two primary questions: (1) Is habitat structural complexity associated with changes in brain size at both the community level and within-species? Drawing from Shumway and Pollen’s interspecies comparison studies on cichlid fish, one might predict that if structural complexity directly imposes greater cognitive demands, then populations from more complex habitats should exhibit larger brains. However, based on our previous findings that fish in less complex habitats perform better in cognitive tests (Jungwirth et al., 2024), we predict that fish from less complex habitats might actually have larger brains. (2) Does variation in habitat structural complexity lead to changes in brain architecture? Given the plasticity of fish brains (Ebbesson & Braithwaite, 2012; Eifert et al., 2015; Sukhum et al., 2018), we hypothesized that different selection might operate separately on different functional regions, leading to their relative enlargement or reduction.

## Methods

### Sampling Habitats

Fieldwork was conducted between April 15 and May 2, 2022, at four distinct habitat sites (hereafter habitats A, B, C, and D) across two islands on Lake Tanganyika, Zambia (**Figure 1**): Chikonde (Mutondwe) Island (8°42′43″ S, 31°05′33″ E) and Mbita (Nkumbula) Island (8°45′15″ S, 31°05′06″ E). See **Table S2** for a summary of habitat traits.

**Figure 1.**
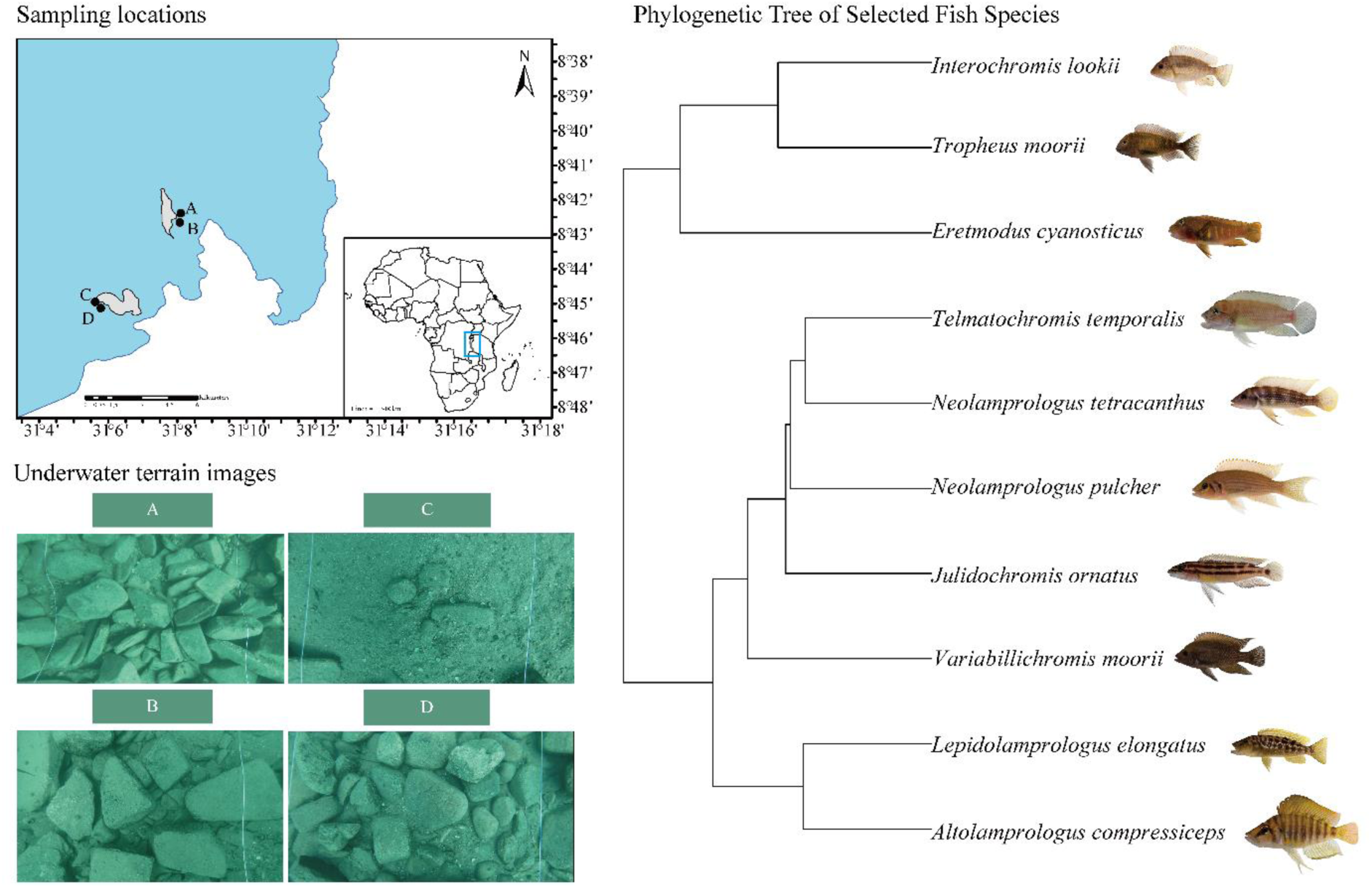
Habitat complexity assessment and sampled species. The map (left) shows the geographic locations of four sampling sites on Lake Tanganyika, Zambia. Mbita Island (8°45′15″ S, 31°05′06″ E) and Chikonde Island (8°42′43″ S, 31°05′33″ E), along with detailed underwater terrain images of each site (Habitat B, Habitat A, Habitat C, Habitat D; top right). The phylogenetic tree (right) illustrates the phylogenetic relationships of the ten sampled species.

### Habitat Selection and Characteristics

Initial habitat selection was conducted by SCUBA divers based on visual assessments of habitat complexity. Habitat A was densely covered with rocks of relatively uniform size, forming a highly complex environment. Habitat D also had a dense rock cover, but the rock sizes varied more significantly, with some large rocks interspersed, resulting in a less uniform but still complex environment. Habitat B was characterized by fewer rocks than Habitats A and D, with a heterogeneous distribution of varying rock sizes, creating clustered formations across the sandy substrate. In contrast, Habitat C had smaller rocks distributed sparsely and evenly, forming a more open and uniform landscape.

### 3D Reconstruction of the Four Habitats

To objectively measure the habitat complexity, a structure-from-motion (SfM) approach (Hartley & Zisserman, 2003) was used to generate 3D reconstructions of the lakebed within a 6 m × 6 m area for each habitat, extending by 0.5 m beyond each grid edge to capture representative structural features (**Figure 2**). Following the protocol described by (Jungwirth et al. 2021), we performed underwater video scanning with a GoPro Hero 5 camera at approximately 0.5 m above the lakebed, ensuring comprehensive coverage. Overlapping video frames were extracted at 1.5 Hz to create high-resolution 3D meshes using Meshroom (version 2023.1.0, Griwodz et al., 2021). These meshes were imported into Blender (version 3.6), where they were cropped to standardized 5 m × 5 m areas for further analysis.

### Complexity Assessment

The structural complexity of each habitat was quantified using three complementary methods: a **rugosity** calculation, **1/Maximum Curvature** (for simplicity, throughout this text, we refer to 1/Max Curvature as **Inverse Curvature**) mapping, and **height distribution** analysis. Together, these methods provided a multi-scale perspective on surface roughness and topographical variation across the habitats.

**Rugosity** was calculated as the ratio of the actual surface area to the projected surface area of a flat plane of the same size, following established protocols (Jungwirth et al., 2024; Pollen et al., 2007a; Shumway, 2008). Higher rugosity values indicate greater surface complexity and unevenness. The rugosity values for each habitat were derived from the 3D reconstructions. Although rugosity provides an overarching measure of surface complexity, it lacks the finer resolution offered by the subsequent curvature and height distribution analyses.

To assess local surface complexity, first, the Maximum Curvature was calculated for each point, representing the principal curvature and indicating the sharpness of surface bends. Then the reciprocal, **Inverse Curvature**, was calculated to reflect the radius of a sphere that would fit the local surface, with larger values corresponding to flatter regions and smaller values indicating higher curvature areas. This metric was selected for its ability to capture fine-scale surface details, complementing the broader measure of rugosity. Probability distribution histograms for Inverse Curvature were created to visualize surface uniformity and the distribution of landmarks across habitats. The standard deviation and maximum values of these distributions represent overall landmark uniformity and localized clustering, respectively. For a comprehensive discussion of the physical relevance and rationale for using Inverse Curvature, see **supplementary material**.

**Figure 2.**
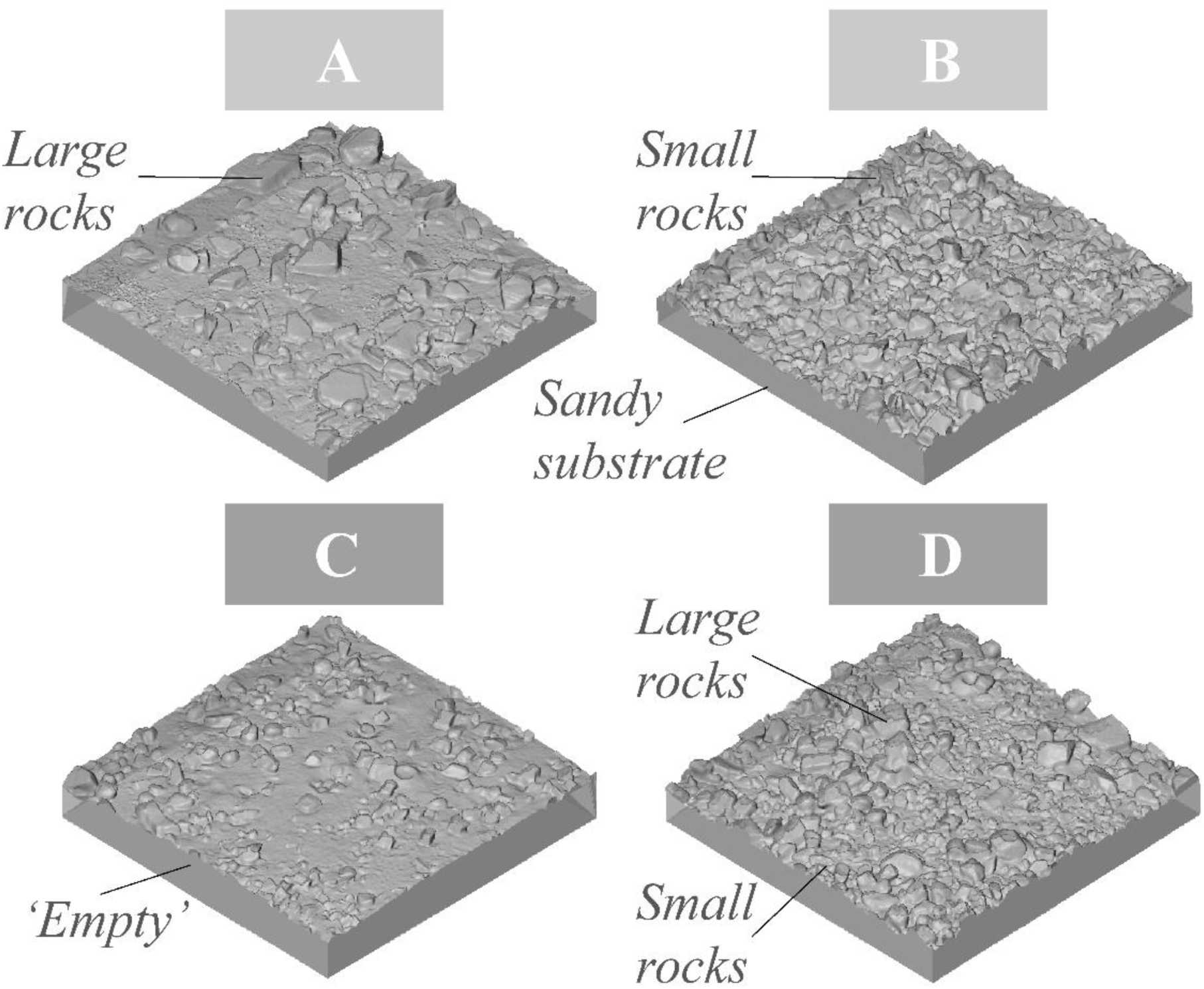
Features of 3D reconstructions of the lake bottom, emphasizing the assessed complexity of topographical features. These sites were selected based on their unique environmental complexities, specifically, Habitat C, due to its sparse rock distribution, was selected as the lowest complexity habitat. The remaining three sites—Habitat D, Habitat A, and Habitat B—represent varying levels of higher complexity, based on the number, size, and distribution of rocks.

**Height distribution** was calculated as the vertical distance between the highest surface points and the base within each habitat. This method captures elevation variation, highlighting the presence or absence of prominent landmarks and aiding in identifying key topographical features. Like the Inverse Curvature maps, height distribution maps offer insights into spatial variation and complement rugosity and curvature metrics. Color-coded maps were produced to facilitate direct comparisons of height distribution across habitats, revealing areas of significant elevation change versus flatter regions.

### Study Species

Ten species of cichlids that occurred at every site were selected for analysis (**Figure 1**). These species occupy varied ecological niches yet show consistent behaviours within each niche. Their territorial behaviour supports reliable data collection and implies long-term residency within the habitats, allowing us to reasonably attribute brain morphology differences to environmental influences. After excluding six samples due to dissection complications, the final dataset comprised 141 fish.

### Data Analysis

To examine the impact of habitat complexity on brain size while controlling for confounding variables, we employed linear mixed-effects models with model comparison and selection. Total brain volume, brain region volumes, and body mass data were log-transformed to meet residual normality assumptions. Body mass was included as a covariate in each model due to its known correlation with brain volume, enabling an isolated evaluation of other variables. Species were treated as a random effect to adjust for intra-species variability, and model fits were evaluated using Akaike Information Criterion (AIC) and Likelihood Ratio Tests (LRT). Models with ΔAIC < 2 were considered equivalent, while those with ΔAIC > 2 were deemed superior (Burnham et al., 2011; Symonds & Moussalli, 2011).

We initially tested ‘habitat’ as a primary factor to determine if brain volume differed significantly across habitats. Based on habitat assessments, habitats of similar complexity were grouped to create a new categorical variable, ‘complexity’, to evaluate any improvement in model performance. To account for potential confounders (e.g., sex and water depth), models were constructed incorporating these covariates separately and in combination with ‘complexity.’ Visibility and water temperature were excluded due to their uniformity across habitats. To evaluate if habitat complexity affected body condition, a mixed-effects model was fitted, relating body mass to standard length and habitat complexity. This model allowed for an assessment of potential differences in energy intake between habitats, as well as the resulting variations in body condition.

#### Species-Specific Analysis

To determine if habitat complexity influenced brain volume variations within species, we conducted linear regression analyses for each species. Using total brain volume as the response variable and habitat complexity (binary: low vs. high; see Results) as the predictor, with body weight as a covariate, we assessed intra-species trends. For visualization, residuals were computed from regression slopes of brain part sizes against body weight for each species.

#### Brain Architecture Analysis

To investigate potential enlargement of specific brain regions, we fitted linear mixed-effects models for each brain region, with habitat complexity as the primary variable and total brain volume as a control covariate. Species were included as a random effect to adjust for species-specific variability in brain architecture.

## Results

### Habitat C is the least complex compared to the other three habitats

By integrating rugosity, Inverse Curvature, and height distribution, we comprehensively assessed habitat complexity. Results indicated that **Habitat C** had the most uniform structure, with minimal elevation changes and lower surface curvature, establishing it as the least complex habitat. Conversely, Habitats B and D displayed higher variability, with elevated rugosity and dispersed landmarks as reflected in both curvature and height distributions.

The **rugosity** values were 1.37 for Habitat A, 1.7 for Habitat B, 1.01 for Habitat C, and 1.7 for Habitat D. This indicates that Habitat C had the overall smoothest surface, while Habitat B, C and D had the most structurally complex surfaces.

The **height distribution** maps (**Figure 2)** corroborated the findings from the curvature analysis. Habitat A had more aggregated smaller rocks, while Habitat B showed local elevation variations with large rock accumulations, and Habitat D had a wider dispersion of rocks with pronounced variations. In comparison, Habitat C demonstrated the most homogeneous height distribution, further confirming that its surface has the least topographical complexity.

**Figure 2.**
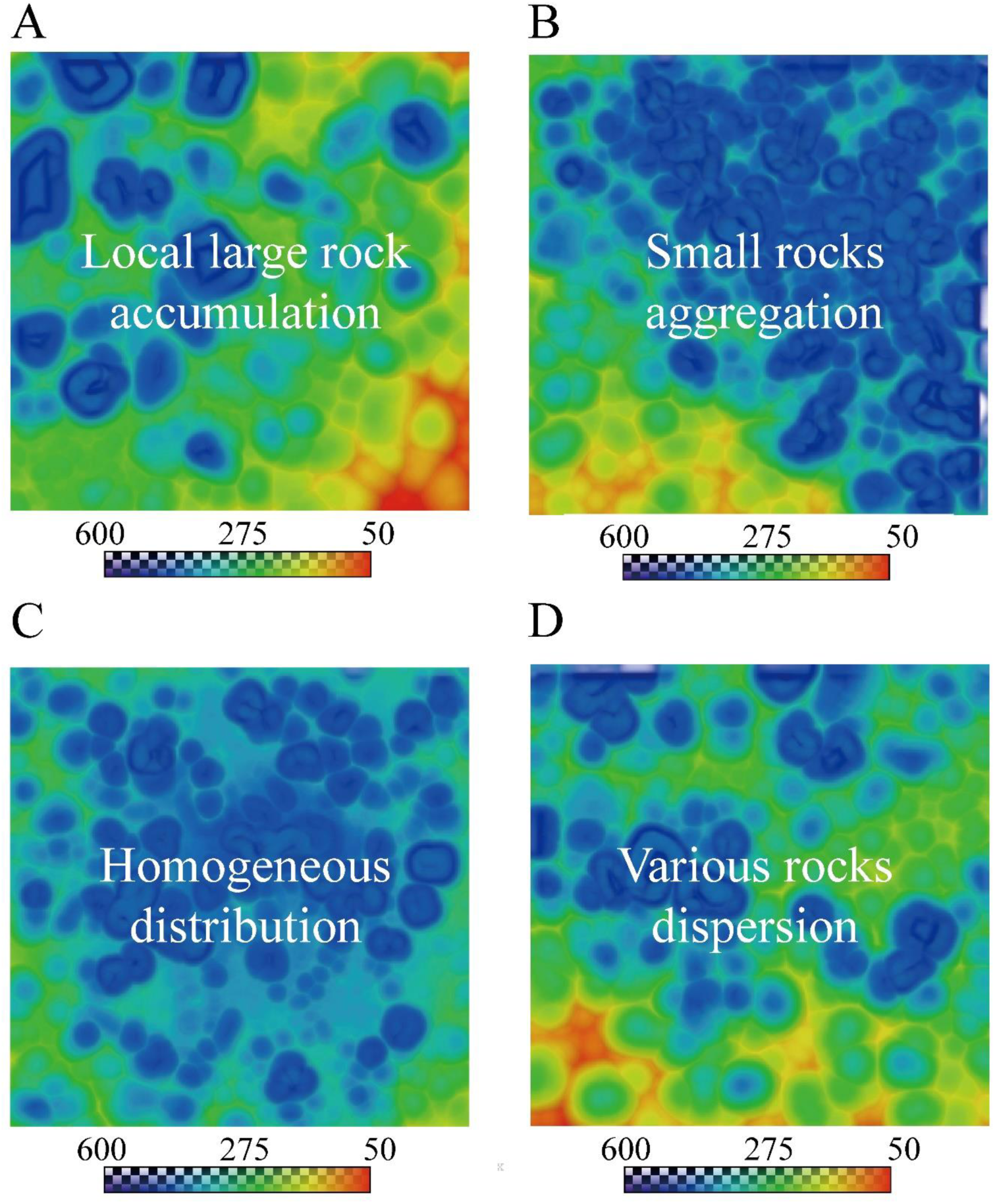
Height Distribution Maps Highlighting Lower Structural Complexity in Habitat C. Warmer colours (yellow/red) indicate regions with lower elevation, while cooler colours (blue/green) represent greater elevation. Habitat B displays elevation variation with large rock accumulations, Habitat A shows smaller rocks aggregated, Habitat C exhibits the most homogeneous surface with minimal elevation variation, and Habitat D displays dispersed rocks with varying heights.

Additionally, the frequency distribution histogram of **Inverse Curvature** for the four habitats (A, B, C, D) is shown in **Figure 3**. This comparison illustrates the varying degrees of terrain complexity, with habitat C being the least complex, while habitats A, B, and D exhibit similar levels of variability.

**Figure 3.**
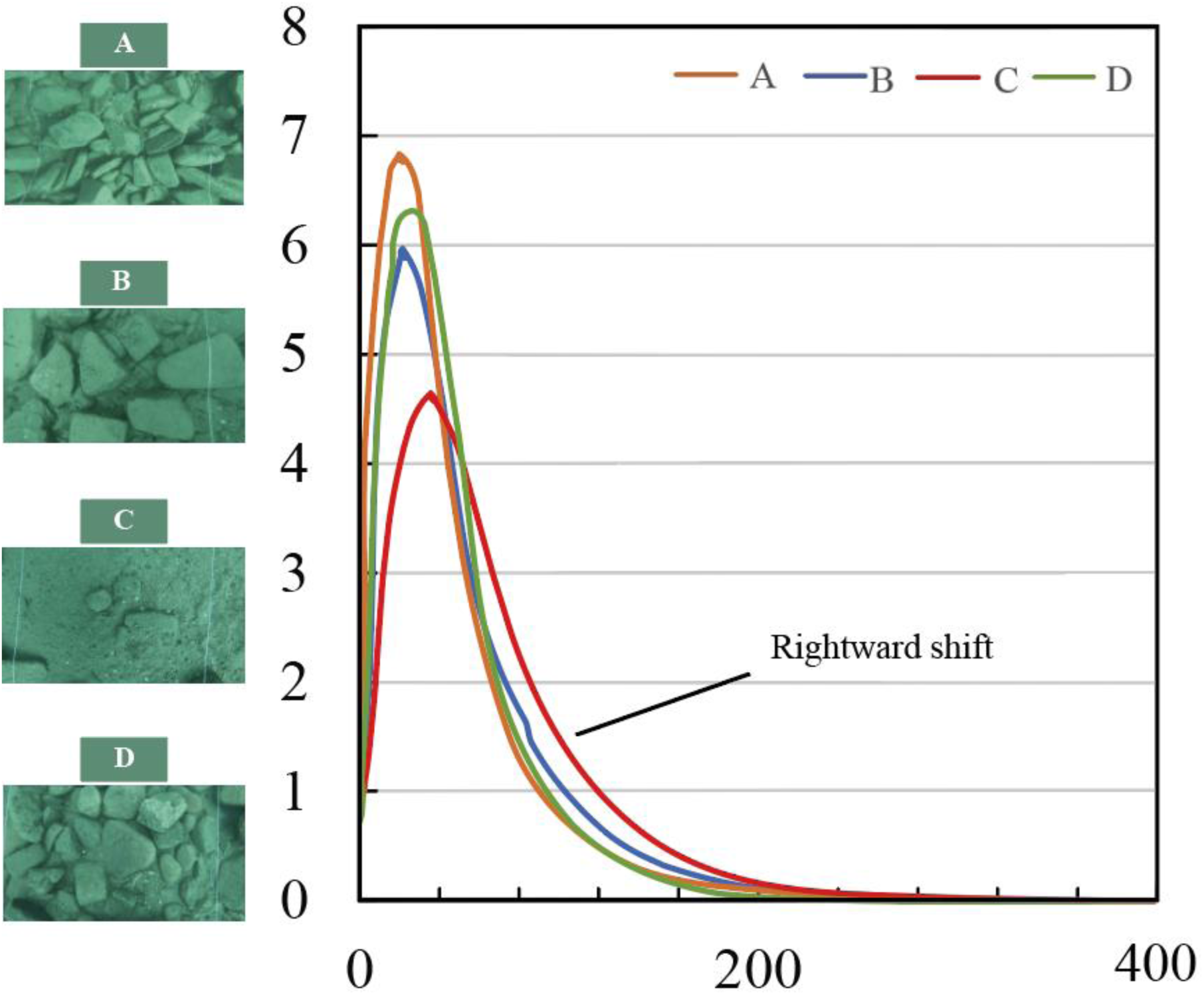
Frequency distribution histogram of Inverse Curvature for four different habitats (A, B, C, D) with corresponding terrain images shown above. The Inverse Curvature values represent surface flatness, where larger values indicate flatter areas, and smaller values indicate more curved or irregular regions. Habitat C shows a notable rightward shift in the distribution compared to the other habitats, suggesting a smoother surface and more uniform terrain. This comparison illustrates the varying degrees of terrain complexity, with habitat C being the least complex, while habitats A, B, and D exhibit similar levels of variability.

### Fish in habitat C have relative larger brains than the other three habitats

Fish populations in the least complex Habitat C were found to have significantly larger brain volumes. **Figure 4** illustrates the relationship between habitat complexity and brain volume. A LMM revealed significant differences in brain volumes across habitats at inter-population level (χ2 = 40.659, df = 3, p < 0.00001). Subsequent Tukey’s HSD tests indicated that fish in Habitat C exhibited significantly larger brain volumes compared to those from Habitat D, Habitat B, and Habitat A. In detail, Habitat C fish had brain volumes 17.59% larger than Habitat D (0.162 ± 0.0402, t = 4.031, df = 128, p = 0.0005), 27.06% greater than Habitat B (0.23952 ± 0.0420, t = 5.708, df = 127, p < 0.0001), and 26.10% larger than Habitat A (0.23188 ± 0.0455, t = 5.096, df = 128, p < 0.0001). No significant differences were found among the other habitats (Tukey’s HSD test, p-values all exceeding 0.24).

**Figure 4.**
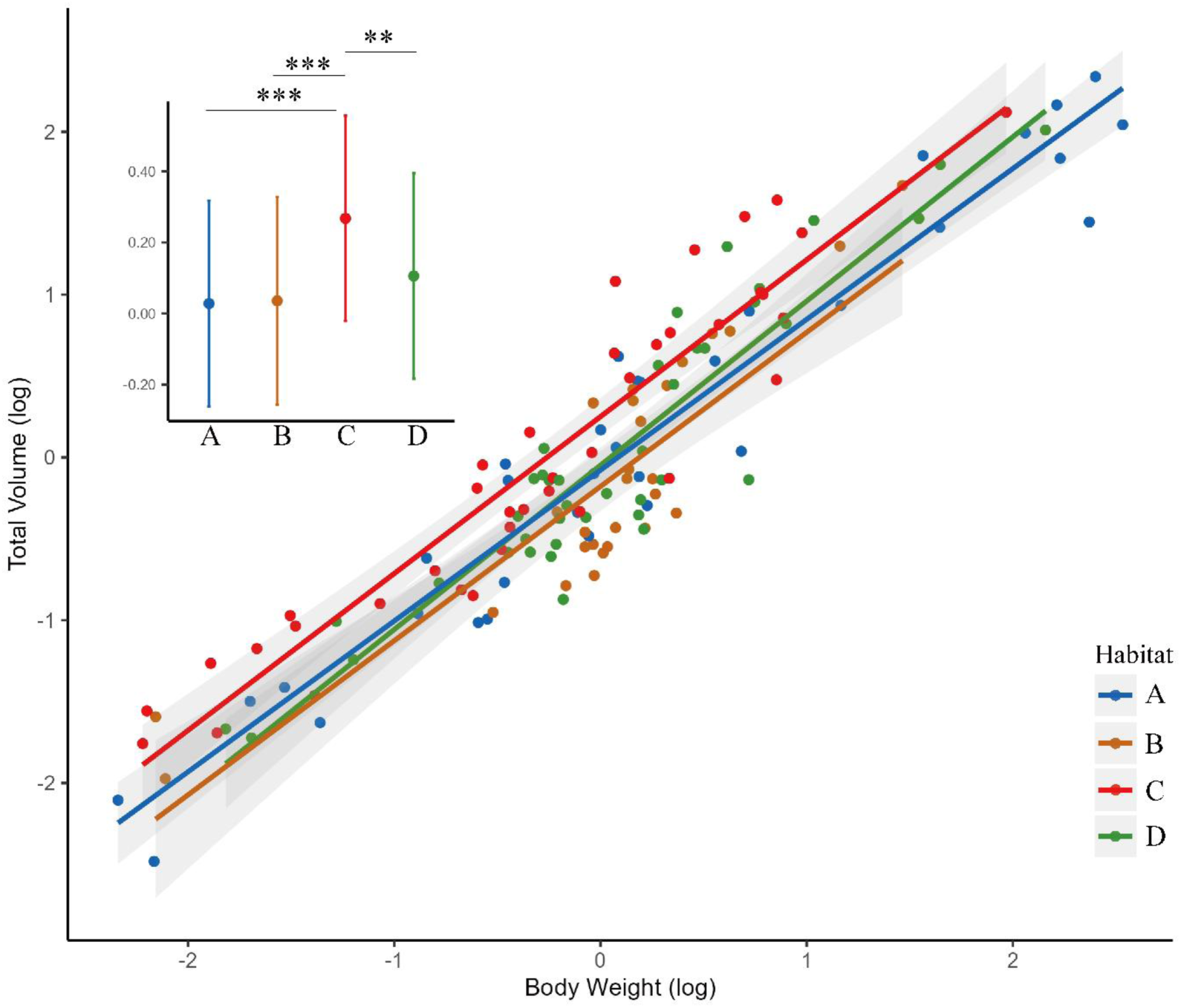
Fish from habitat C have relatively larger brains. Regression lines and 95% confidence intervals depict the total volume of the brain transformed (mm^3^) as a function of the transformed body weight (mg) for each population: Habitat A (blue), Habitat B (orange), Habitat C (red), and Habitat D (green). The inset displays estimated marginal means for each population with 95% confidence intervals derived from the statistical models. Significance markers denote comparisons where Habitat C shows a statistically significant larger brain volume compared to Habitat A (***P < 0.001), Habitat B (***P < 0.001), and Habitat D (**P < 0.01): LMM followed by Tukey’s HSD test.

Larger brains in low complexity habitats

To enhance statistical power, we grouped habitats A, B, and D, which exhibit similar structural complexity, into a new category labeled as **’high complexity’**. Habitat C, characterized by the lower structural complexity, was categorized as **’low complexity’**. This new categorical variable, **’complexity’**, was then used as a predictor in the subsequent analyses.

We found a consistent effect of habitat complexity on brain volume across species. Our regression analysis, as detailed in **Table S3**. and illustrated in **Figure 5**, revealed a uniform trend across species where populations from habitats of ‘low complexity’ consistently exhibited larger brain volumes. Although not all species showed statistically significant differences, this pattern supports the hypothesis that lower environmental complexity correlates with increased brain volume within species. In particular, *Julidochromis ornatus*, *Neolamprologus tetracanthus*, *Telmatochromis temporalis*, and *Tropheus moorii* demonstrated significantly larger brain volumes in less complex environments (p < 0.05), highlighting their sensitivity to habitat complexity.

**Figure 5.**
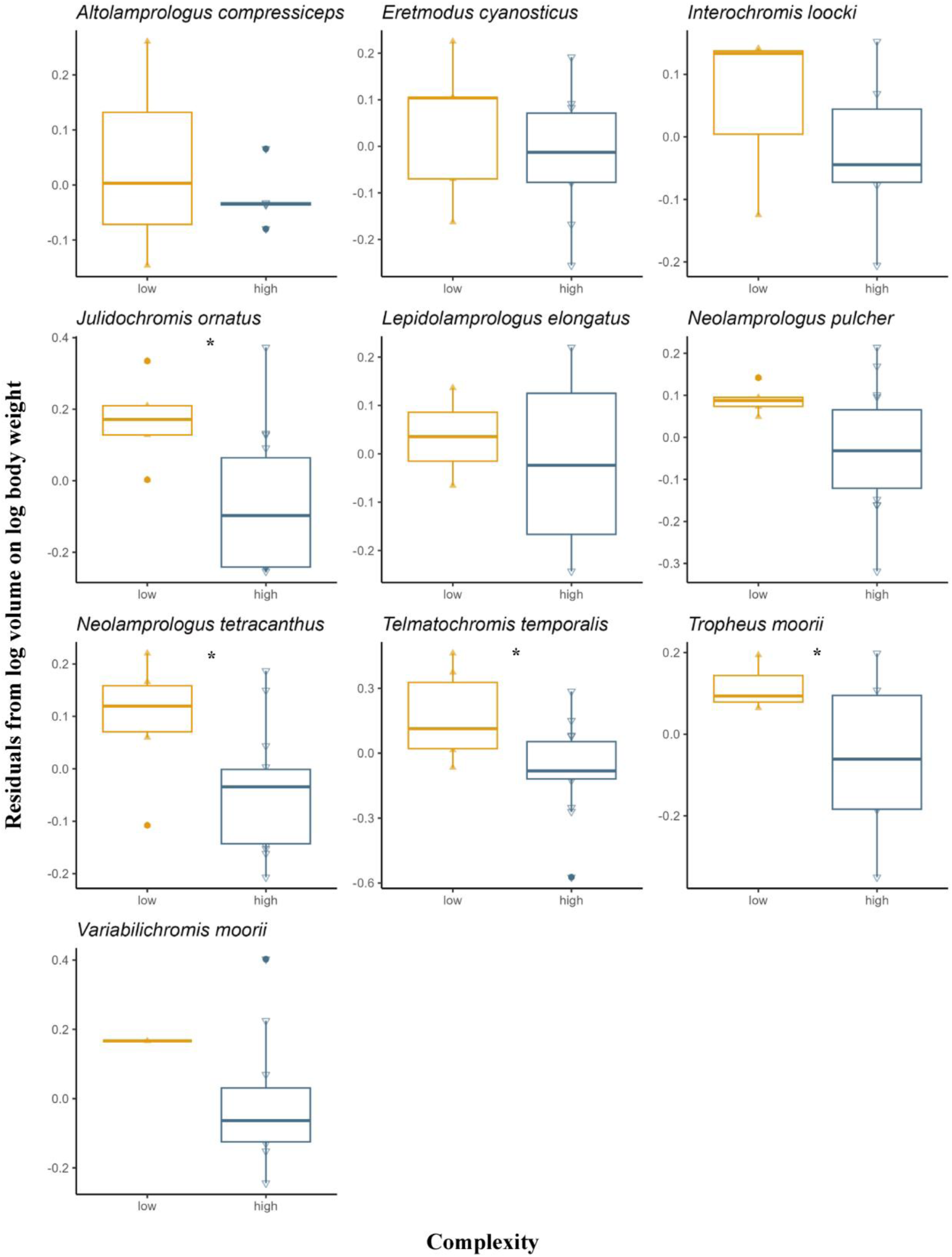
Larger brain volumes in low complexity habitats. Box plots representing the residuals of brain size (adjusted for body weight) against habitat complexity for ten species. Each plot contrasts the residuals for species in high- and low-complexity environments, illustrating the variation within and across the groups.

### Differential brain region responses to habitat complexity: cerebellum enlargement and hypothalamus reduction

For brain architecture, we observed a significant enlargement of the cerebellum in fish from low-complexity habitat (Habitat C; see **Figure 6**), with an increase of 12.56% relative to the rest of the brain (β = 0.118322, t (130.448) = 2.082, p = 0.0393), suggesting enhanced adaptability to environment structure. In contrast, the hypothalamus exhibited a non-significant trend towards reduction by 11.09% in the less complex habitat (Habitat C) (β = −0.11753, t (133.41) = −1.637, p = 0.104). No significant size changes were found in other brain areas, indicating a more conserved developmental pattern across varying levels of habitat complexity. These findings underscore the unique evolutionary response of the cerebellum (and hypothalamus), supporting the hypothesis that brain regions may evolve independently to meet specific ecological demands.

**Figure 6.**
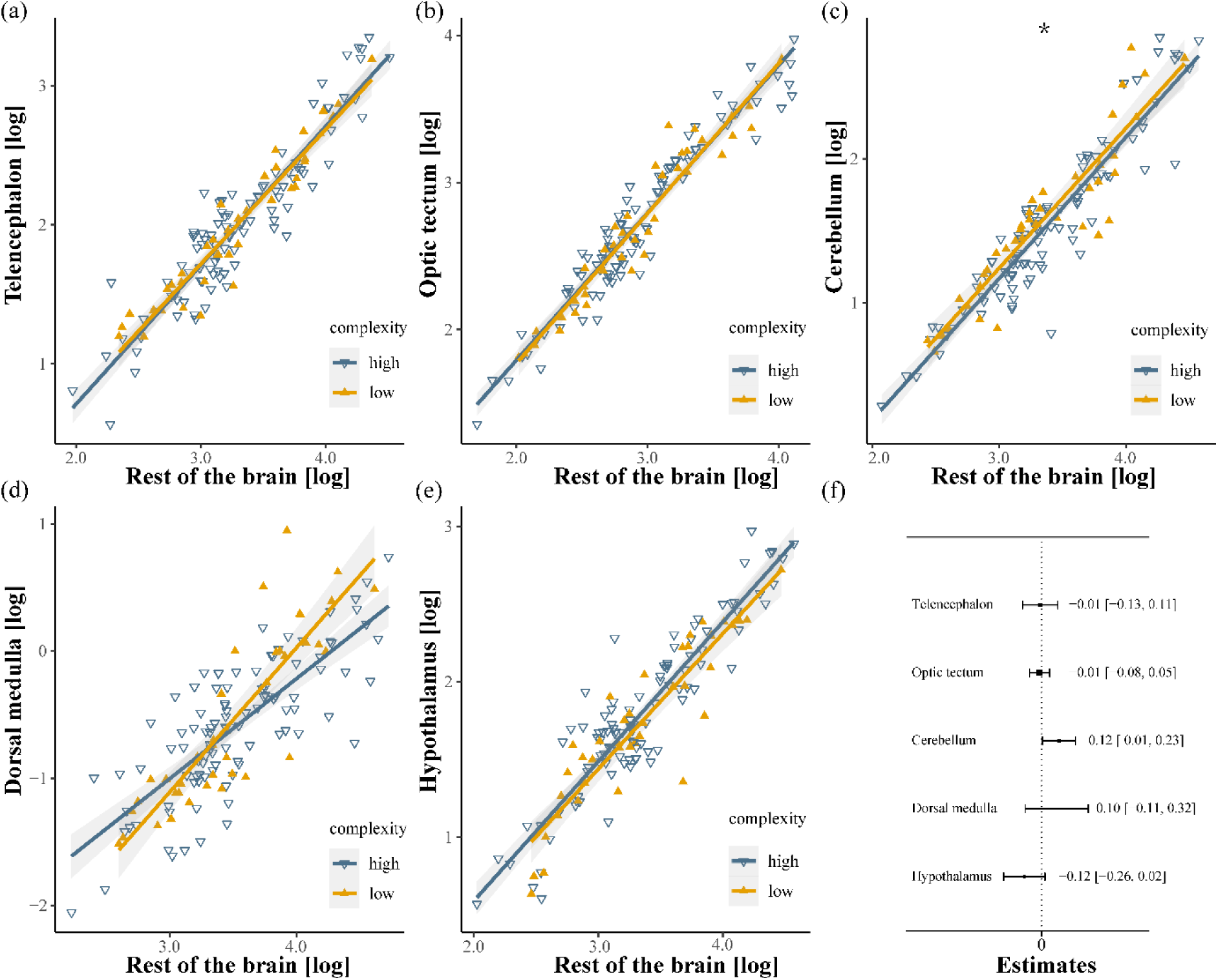
Relative brain region size between habitat complexity. This figure shows regression analyses for five regions of the brain: (a) Telencephalon, (b) Optic Tectum, (c) Cerebellum, (d) Dorsal Medulla and (e) Hypothalamus, each plotted against the rest of the size of the brain on a logarithmic scale. Panel (f) summarises the effect size estimates with confidence intervals for each region. In particular, only the cerebellum shows a significant enlargement in low complexity environments, with an estimate of 0.12 and a confidence interval of [0.01, 0.23] (*P < 0.05).

## Discussion

By comparing ten species of Tanganyikan cichlids across the same four habitats, we find that habitat structural complexity can influence brain size at both community and species level. In contrast to previous work comparing among species, we observed a negative association between habitat complexity and brain size; specifically, fish from the less complex habitat (C) exhibited significantly larger brain volumes than those from more complex habitats (A, B, D), with increases of 27.06%, and 26.10%, and 17.59% respectively. This pattern was consistent across species, with statistically significant results in four out of ten species and a similar trend in the remaining six. Additionally, our analysis revealed distinctive changes in brain architecture: in less complex habitats, the cerebellum was disproportionately larger (cerebellum to rest of brain) by 12.56% compared to more complex habitats, whereas the hypothalamus tended to decrease in proportion. Together with previous research, which demonstrated that fish from the less complex areas exhibited better cognitive performance (Jungwirth et al., 2024), our results do not support the clever foraging hypothesis (CFH; George F. Striedter et al., 2005; P. Park & Bell, 2010). Because our design allows us to compare both among and within species across the same habitats, we suggest that each species may experience relationship between habitat structural complexity and cognitive demand in different ways.

### Challenges in testing CFH

There are two main challenges in testing the CFH. First, most studies exploring the relationship between brain morphology and habitat use rely on interspecific comparisons, while intraspecific comparisons conducted in the wild are relatively scarce. Yet interspecies comparisons risk conflating current functionality with historical adaptations (Harvey et al., 1980; Healy & Rowe, 2007; Huber et al., 1997; Pollen et al., 2007; Powell & Leal, 2014; Shumway, 2008) and although interspecific comparisons certainly offer valuable insights due to the significant ecological divergence between species, intraspecific comparisons are essential for isolating the specific effects of habitat use on brain morphology (Gonda et al., 2013). It is also noteworthy that intraspecific comparisons have yielded conflicting results for the CFH. For example, in pumpkinseed sunfish, fish from more structurally complex littoral ecotypes had larger overall brain sizes (Caleb J. Axelrod et al., 2018)., while in three-spine stickleback there was no consistent change in brain morphology with habitat complexity (Ahmed et al., 2017).

Another challenge lies in our ability to quantify environmental complexity. Despite the increasing attention to environmental complexity, our methods for quantifying the habitat structural patterns remain limited. The habitats selected in previous studies are often categorized by their broad ecotypes or ecological classifications like sandy, rocky or vegetated habitats. This simplification can be confounded by small scale variation that the animals actually experience, thereby complicating the interpretation of research results. To address this issue, researchers have employed quantitative measurement such as "rugosity" and “optical intensity” to assess complexity (Shumway et al., 2007), but studies that adopt these methods still overlook more nuanced physical characteristics like substrate homogeneity (González-Rivero et al., 2017; Gratwicke & Speight, 2005; Loke & Chisholm, 2022; Powell & Leal, 2014; Shumway, 2008; Storks et al., 2024). Therefore, to deepen our understanding of how habitat structural complexity affects cognitive performance and brain morphology, further studies that use more detailed, quantified measures of structural complexity across habitats within the same ecotype are essential.

### Cognitive Demands in Low Complexity Environments

While numerous studies indicate that higher environmental complexity correlates with increased brain size (DePasquale et al., 2016; di Porzio, 2020; Pollen et al., 2007; Safi & Dechmann, 2005; Shumway, 2008, 2010; Sobrero et al., 2016; White & Brown, 2015), our research is among the first to identify a negative correlation between low structural complexity and larger brain volumes within populations of the same ecotype but inhabiting different environments. Why might lower complexity correlate with larger brains? In the fish populations we studied, environments with lower complexity could impose higher cognitive demands, such as spatial memory and navigation. A recent study has shown that in environments with low habitat complexity, Western mosquitofish exhibit improved cognition, attributed to increased risk-taking behaviour. This difference in cognition is likely driven by the scarcity of refuges and heightened predation pressure (Irwin et al., 2024). This hypothesis closely aligns with the unique behaviours of our study species, contrasting with findings from research on Lake Tanganyika’s Ectodine clade cichlids, which showed that species originally caught in complex, rocky habitats have larger brain volumes (Pollen et al., 2007). Our study used seven Lamprologines, territorial species known for their extensive ecological niches and rapid adaptation across diverse environments (Konings, 2019). These species primarily inhabit and reproduce in rock crevices, relying on rocks as essential resources, and a reduction in the overall abundance of refuges could increase the cognitive demands of finding suitable shelter, consistent with existing research demonstrating that harsh environments are associated with larger brain sizes (Howell & Walsh, 2023; Roth & Pravosudov, 2009; Walsh et al., 2016). In our study, the low complexity of the Habitat C is reflected in two main aspects and both could generate greater cognitive challenges. First, the rocks in Habitat C are fewer and smaller, limiting the available hiding spots for the fish, which could expose them to greater predation pressure known to be associated with an increase in brain size in both wild (Walsh et al., 2016) and laboratory studies (Reddon et al., 2018), as well with increased foraging difficulty, also known to increase brain size (e.g. Edmunds et al., 2016). Second, the substrate distribution in Habitat C is highly uniform, with few prominent landmarks and geometric information (as detailed in the results section). Studies have shown that cichlids rely on landmarks and geometric information for navigation (Brown et al., 2007; Suriyampola & Eason, 2015), and the absence of such landmarks requires them to engage in more precise spatial memory, thus increasing cognitive challenges.

### Changes in Brain Structure

In this study, we also investigated whether environmental complexity could exert differential pressures on specific regions of the brain, leading to structural changes. The results showed that the cerebellum was disproportionately enlarged, potentially reflecting the specific selective pressures faced by the fish in low complexity habitat. In the low-complexity Habitat C, limited hiding spots, increased predation difficulty, and greater risk of predation not only challenge general cognition but also impose higher demands on navigation and motor skills. The cerebellum has been well documented involved in spatial navigation and well-known in motor control (Durán et al., 2014; Martin et al., 2003; Mirino et al., 2022; Rochefort et al., 2013). Moreover, recent studies on mosquitofish have similarly found that individuals in low-complexity environments not only completed spatial cognitive challenges more quickly but also exhibited stronger motor control capabilities (Irwin et al., 2024).

It is important to note that our study does not identify the proximate mechanisms underlying the observed association between habitat complexity and brain size. Three potential mechanisms could explain the variation in brain size across different ecotypes within varied habitats: habitat choice, phenotypic plasticity, and diversifying selection. Firstly, individuals may select habitats that match a brain phenotype optimized for specific environmental conditions (Edelaar & Bolnick, 2019). Secondly, brain size and morphology might adapt over an individual’s lifetime if phenotypic plasticity allows adjustments to local environmental conditions (Eifert et al., 2015; Gonda et al., 2009; McCallum et al., 2014). Lastly, different cognitive and ecological demands across habitats could drive selection pressures that favour divergent brain morphologies, as evidenced by experiments showing artificial selection for brain size in guppies (*Poecilia reticulata*), suggesting heritable changes under sustained selection (A. Kotrschal et al., 2013). However, our comparative study design cannot determine which mechanism, or combination of mechanisms, is responsible. Further investigation using reciprocal transplant or common garden experiments, like those conducted by (Walsh et al., 2016), would be needed to clarify these mechanisms.

Our study, using ten species across four habitats provides community and within-species evidence of a more nuanced relationship between habitat structural complexity and brain size, with lower habitat complexity associated with larger brain sizes and distinct architecture changes. This suggests that simple environment may impose sophisticated cognitive and ecological demands.

## Acknowledgements

We thank the Department of Fisheries in Mpulungu, Zambia for kindly supporting our research at Lake Tanganyika. We are grateful to the staff at the Tanganyika Science Lodge and Kalambo Falls Lodge for their hospitality.

## Contribution Statement

**Bin Ma:** Conceptualization (lead); data collection and curation (lead); formal analysis (lead); investigation (lead); methodology (lead); project administration (equal); resources (equal); visualization (lead); writing – original draft (lead); writing – review and editing (lead). **Arne Jungwirth:** Conceptualization (co-lead); Data collection (lead); writing – review and editing (supporting); resources (supporting). **Weiwei Li:** data collection (supporting); writing – review and editing (supporting). **Zitan Song:** formal analysis (supporting); writing – review and editing (supporting). **Stefan Fischer:** data collection (supporting); writing – review and editing (supporting). **Etienne Lein:** data collection (supporting); writing – review and editing (supporting). **Alex Jordan:** Supervision (lead); Funding acquisition (lead); project administration (lead); resources (lead); writing – review and editing (co-lead).

## Data Accessibility

All associated data are available at DRYAD

## References

Ahmed, N. I., Thompson, C., Bolnick, D. I., & Stuart, Y. E. (2017). Brain morphology of the threespine stickleback (Gasterosteus aculeatus) varies inconsistently with respect to habitat complexity: A test of the Clever Foraging Hypothesis. Ecology and Evolution, 7(10), 3372–3380. 10.1002/ece3.2918

Benson-Amram, S., Dantzer, B., Stricker, G., Swanson, E. M., & Holekamp, K. E. (2016). Brain size predicts problem-solving ability in mammalian carnivores. Proceedings of the National Academy of Sciences of the United States of America, 113(9), 2532–2537. 10.1073/pnas.1505913113

Brown, A. A., Spetch, M. L., & Hurd, P. L. (2007). Growing in Circles: Rearing Environment Alters Spatial Navigation in Fish: (603982013-031) [Dataset]. 10.1037/e603982013-031

Bshary, R., Wickler, W., & Fricke, H. (2002). Fish cognition: A primate’s eye view. Animal Cognition, 5(1), 1–13. 10.1007/s10071-001-0116-5

Caleb J. Axelrod, Caleb J. Axelrod, Frédéric Laberge, Frederic Laberge, Beren W. Robinson, & Beren W. Robinson. (2018). Intraspecific brain size variation between coexisting sunfish ecotypes. Proceedings of The Royal Society B: Biological Sciences. 10.1098/rspb.2018.1971

Chittka, L., & Niven, J. (2009). Are Bigger Brains Better? Current Biology, 19(21), R995–R1008. 10.1016/j.cub.2009.08.023

Deaner, R. O., Isler, K., Burkart, J., & van Schaik, C. (2007). Overall brain size, and not encephalization quotient, best predicts cognitive ability across non-human primates. Brain, Behavior and Evolution, 70(2), 115–124. 10.1159/000102973

DePasquale, C., Neuberger, T., Hirrlinger, A. M., & Braithwaite, V. A. (2016). The influence of complex and threatening environments in early life on brain size and behaviour. Proceedings of the Royal Society B: Biological Sciences, 283(1823), 20152564. 10.1098/rspb.2015.2564

di Porzio, U. (2020). A bigger brain for a more complex environment. Reviews in the Neurosciences, /j/revneuro.ahead-of-print/revneuro-2020–0041/revneuro-2020-0041.xml. 10.1515/revneuro-2020-0041

Dobberfuhl, A. P., Ullmann, J. F. P., & Shumway, C. A. (2005). Visual acuity, environmental complexity, and social organization in african cichlid fishes. Behavioral Neuroscience, 119(6), 1648–1655. 10.1037/0735-7044.119.6.1648

Durán, E., Ocaña, F. M., Martín-Monzón, I., Rodríguez, F., & Salas, C. (2014). Cerebellum and spatial cognition in goldfish. Behavioural Brain Research, 259, 1–8. 10.1016/j.bbr.2013.10.039

Ebbesson, L. O. E., & Braithwaite, V. A. (2012). Environmental effects on fish neural plasticity and cognition. Journal of Fish Biology, 81(7), 2151–2174. 10.1111/j.1095-8649.2012.03486.x

Edelaar, P., & Bolnick, D. I. (2019). Appreciating the Multiple Processes Increasing Individual or Population Fitness. Trends in Ecology & Evolution, 34(5), 435–446. 10.1016/j.tree.2019.02.001

Edmunds, N. B., Laberge, F., & McCann, K. S. (2016). A role for brain size and cognition in food webs. Ecology Letters, 19(8), 948–955. 10.1111/ele.12633

Eifert, C., Farnworth, M., Schulz-Mirbach, T., Riesch, R., Bierbach, D., Klaus, S., Wurster, A., Tobler, M., Streit, B., Indy, J. R., Arias-Rodriguez, L., & Plath, M. (2015). Brain size variation in extremophile fish: Local adaptation versus phenotypic plasticity. Journal of Zoology, 295(2), 143–153. 10.1111/jzo.12190

Gemma E. White, Gemma E. White, Culum Brown, & Culum Brown. (2015). Microhabitat Use Affects Brain Size and Structure in Intertidal Gobies. Brain Behavior and Evolution. 10.1159/000380875

George F. Striedter, Georg F. Striedter, & George F. Striedter. (2005). Principles of brain evolution.

Gonda, A., Herczeg, G., & Merilä, J. (2009). Adaptive brain size divergence in nine-spined sticklebacks ( *Pungitius pungitius*)? Journal of Evolutionary Biology, 22(8), 1721–1726. 10.1111/j.1420-9101.2009.01782.x

Gonda, A., Herczeg, G., & Merilä, J. (2013). Evolutionary ecology of intraspecific brain size variation: A review. Ecology and Evolution, 3(8), 2751–2764. 10.1002/ece3.627

González-Rivero, M., Harborne, A. R., Herrera-Reveles, A., Bozec, Y.-M., Rogers, A., Friedman, A., Ganase, A., & Hoegh-Guldberg, O. (2017). Linking fishes to multiple metrics of coral reef structural complexity using three-dimensional technology. Scientific Reports, 7(1), 13965. 10.1038/s41598-017-14272-5

Gonzalez-Voyer, A., Winberg, S., & Kolm, N. (2008). Social fishes and single mothers: Brain evolution in African cichlids. Proceedings of the Royal Society B: Biological Sciences, 276(1654), 161–167. 10.1098/rspb.2008.0979

Gratwicke, B., & Speight, M. R. (2005). The relationship between fish species richness, abundance and habitat complexity in a range of shallow tropical marine habitats. Journal of Fish Biology, 66(3), 650–667. 10.1111/j.0022-1112.2005.00629.x

Harvey, P. H., Clutton-Brock, T. H., & Mace, G. M. (1980). Brain size and ecology in small mammals and primates. Proceedings of the National Academy of Sciences, 77(7), 4387–4389. 10.1073/pnas.77.7.4387

Healy, S. D., & Rowe, C. (2007). A critique of comparative studies of brain size. Proceedings of the Royal Society B: Biological Sciences, 274(1609), 453–464. 10.1098/rspb.2006.3748

Howell, K. J., & Walsh, M. R. (2023). Transplant experiments demonstrate that larger brains are favoured in high-competition environments in Trinidadian killifish. Ecology Letters, 26(1), 53–62. 10.1111/ele.14133

Huber, R., Van Staaden, M. J., Kaufman, L. S., & Liem, K. F. (1997). Microhabitat Use, Trophic Patterns, and the Evolution of Brain Structure in African Cichlids. *Brain*, Behavior and Evolution, 50(3), 167–182. 10.1159/000113330

Iglesias, T. L., Dornburg, A., Warren, D. L., Wainwright, P. C., Schmitz, L., & Economo, E. P. (2018). Eyes Wide Shut: The impact of dim-light vision on neural investment in marine teleosts. Journal of Evolutionary Biology, 31(8), 1082–1092. 10.1111/jeb.13299

Irwin, K., Aspbury, A. S., Bonner, T., & Gabor, C. R. (2024). Habitat structural complexity predicts cognitive performance and behaviour in western mosquitofish. Biology Letters, 20(7), 20230394. 10.1098/rsbl.2023.0394

Jerison, H. J. (Ed.). (1975). Evolution of the Brain and Intelligence. Current Anthropology, 16(3), 403–426.

Jungwirth, A., Horsfield, A., Nührenberg, P., & Fischer, S. (2024). Estimating Cognitive Ability in the Wild: Validation of a Detour Test Paradigm Using a Cichlid Fish (Neolamprologus pulcher). Fishes, 9(2), Article 2. 10.3390/fishes9020050

Jungwirth, A., Nührenberg, P., & Jordan, A. (2021). On the importance of defendable resources for social evolution: Applying new techniques to a long-standing question. Ethology, 127(10), 872 – 885. 10.1111/eth.13143

Konings, A. (2019). Tanganyika cichlids in their natural habitat (4th Edition).

Kotrschal, A., Rogell, B., Bundsen, A., Svensson, B., Zajitschek, S., Brännström, I., Immler, S., Maklakov, A. A., & Kolm, N. (2013). Artificial selection on relative brain size in the guppy reveals costs and benefits of evolving a larger brain. Current Biology, 23(2), 168–171. Scopus. 10.1016/j.cub.2012.11.058

Kotrschal, K., Van Staaden, M. J., & Huber, R. (1998). Fish Brains: Evolution and Anvironmental Relationships. Reviews in Fish Biology and Fisheries, 8(4), 373–408. 10.1023/A:1008839605380

Loke, L. H. L., & Chisholm, R. A. (2022). Measuring habitat complexity and spatial heterogeneity in ecology. Ecology Letters, 25(10), 2269–2288. 10.1111/ele.14084

MacLean, E. L., Hare, B., Nunn, C. L., Addessi, E., Amici, F., Anderson, R. C., Aureli, F., Baker, J. M., Bania, A. E., Barnard, A. M., Boogert, N. J., Brannon, E. M., Bray, E. E., Bray, J., Brent, L. J. N., Burkart, J. M., Call, J., Cantlon, J. F., Cheke, L. G.,… Zhao, Y. (2014). The evolution of self-control. Proceedings of the National Academy of Sciences, 111(20), E2140–E2148. 10.1073/pnas.1323533111

Martin, L. A., Goldowitz, D., & Mittleman, G. (2003). The cerebellum and spatial ability: Dissection of motor and cognitive components with a mouse model system. European Journal of Neuroscience, 18(7), 2002–2010. 10.1046/j.1460-9568.2003.02921.x

McCallum, E., Capelle, P., & Balshine, S. (2014). Seasonal plasticity in telencephalon mass of a benthic fish. JOURNAL OF FISH BIOLOGY, 85(5), 1785–1792. 10.1111/jfb.12507

Mirino, P., Pecchinenda, A., Boccia, M., Capirchio, A., D’Antonio, F., & Guariglia, C. (2022). Cerebellum-Cortical Interaction in Spatial Navigation and Its Alteration in Dementias. Brain Sciences, 12(5), 523. 10.3390/brainsci12050523

Northmore, D. (2011). The Optic Tectum (pp. 131–142).

Park, P., & Bell, M. (2010). Variation of telencephalon morphology of the threespine stickleback (Gasterosteus aculeatus) in relation to inferred ecology. JOURNAL OF EVOLUTIONARY BIOLOGY, 23(6), 1261–1277. 10.1111/j.1420-9101.2010.01987.x

Park, P. J., & Bell, M. A. (2010). Variation of telencephalon morphology of the threespine stickleback (Gasterosteus aculeatus) in relation to inferred ecology. Journal of Evolutionary Biology, 23(6), 1261–1277. 10.1111/j.1420-9101.2010.01987.x

Pollen, A. A., Dobberfuhl, A. P., Scace, J., Igulu, M. M., Renn, S. C. P., Shumway, C. A., & Hofmann, H. A. (2007). Environmental Complexity and Social Organization Sculpt the Brain in Lake Tanganyikan Cichlid Fish. Brain, Behavior and Evolution, 70(1), 21–39. 10.1159/000101067

Powell, B. J., & Leal, M. (2014). Brain Organization and Habitat Complexity in *Anolis* Lizards. Brain, Behavior and Evolution, 84(1), 8–18. 10.1159/000362197

Reader, S. M., Hager, Y., & Laland, K. N. (2011). The evolution of primate general and cultural intelligence. Philosophical Transactions of the Royal Society B: Biological Sciences, 366(1567), 1017–1027. 10.1098/rstb.2010.0342

Reddon, A. R., Chouinard-Thuly, L., Leris, I., & Reader, S. M. (2018). Wild and laboratory exposure to cues of predation risk increases relative brain mass in male guppies. Functional Ecology, 32(7), 1847–1856. 10.1111/1365-2435.13128

Rochefort, C., Lefort, J., & Rondi-Reig, L. (2013). The cerebellum: A new key structure in the navigation system. Frontiers in Neural Circuits, 7. 10.3389/fncir.2013.00035

Rosati, A. G. (2017). Foraging Cognition: Reviving the Ecological Intelligence Hypothesis. Trends in Cognitive Sciences, 21(9), 691–702. 10.1016/j.tics.2017.05.011

Roth, T. C., & Pravosudov, V. V. (2009). Hippocampal volumes and neuron numbers increase along a gradient of environmental harshness: A large-scale comparison. Proceedings of the Royal Society B: Biological Sciences, 276(1656), 401–405. 10.1098/rspb.2008.1184

Safi, K., & Dechmann, D. K. N. (2005). Adaptation of brain regions to habitat complexity: A comparative analysis in bats (Chiroptera). Proceedings of the Royal Society B: Biological Sciences, 272(1559), 179–186. 10.1098/rspb.2004.2924

Shumway, C. A. (2008). Habitat Complexity, Brain, and Behavior. Brain, Behavior and Evolution, 72(2), 123–134. 10.1159/000151472

Shumway, C. A. (2010). The evolution of complex brains and behaviors in African cichlid fishes. Current Zoology, 56(1), 144–156. 10.1093/czoolo/56.1.144

Shumway, C. A., Hofmann, H. A., & Dobberfuhl, A. P. (2007). Quantifying habitat complexity in aquatic ecosystems. Freshwater Biology, 52(6), 1065–1076. 10.1111/j.1365-2427.2007.01754.x

Sobrero, R., Fernández-Aburto, P., Ly-Prieto, Á., Delgado, S. E., Mpodozis, J., & Ebensperger, L. A. (2016). Effects of Habitat and Social Complexity on Brain Size, Brain Asymmetry and Dentate Gyrus Morphology in Two Octodontid Rodents. Brain Behavior and Evolution, 87(1), 51–64. 10.1159/000444741

Sol, D. (2008). Revisiting the cognitive buffer hypothesis for the evolution of large brains. Biology Letters, 5(1), 130–133. 10.1098/rsbl.2008.0621

Sol, D., Duncan, R. P., Blackburn, T. M., Cassey, P., & Lefebvre, L. (2005). Big brains, enhanced cognition, and response of birds to novel environments. Proceedings of the National Academy of Sciences of the United States of America, 102(15), 5460–5465. 10.1073/pnas.0408145102

Storks, L., Garcia, J., Perez-Martinez, C. A., & Leal, M. (2024). Habitat complexity influences neuron number in six species of Puerto Rican *Anolis*. Biology Letters, 20(2), 20230419. 10.1098/rsbl.2023.0419

Sukhum, K. V., Shen, J., & Carlson, B. A. (2018). Extreme Enlargement of the Cerebellum in a Clade of Teleost Fishes that Evolved a Novel Active Sensory System. Current Biology, 28(23), 3857–3863.e3. 10.1016/j.cub.2018.10.038

Suriyampola, P. S., & Eason, P. K. (2015). The Effects of Landmarks on Territorial Behavior in a Convict Cichlid, *Amatitlania siquia*. Ethology, 121(8), 785–792. 10.1111/eth.12393

Walsh, M. R., Broyles, W., Beston, S. M., & Munch, S. B. (2016). Predator-driven brain size evolution in natural populations of Trinidadian killifish ( *Rivulus hartii* ). Proceedings of the Royal Society B: Biological Sciences, 283(1834), 20161075. 10.1098/rspb.2016.1075

White, G. E., & Brown, C. (2015). Microhabitat Use Affects Brain Size and Structure in Intertidal Gobies. Brain Behavior and Evolution, 85(2), 107–116. 10.1159/000380875

Yopak, K. E., & Lisney, T. J. (2012). Allometric Scaling of the Optic Tectum in Cartilaginous Fishes. Brain, Behavior and Evolution, 80(2), 108–126. 10.1159/000339875

Závorka, L., Koene, J. P., Armstrong, T. A., Fehlinger, L., & Adams, C. E. (2022). Differences in brain morphology of brown trout across stream, lake, and hatchery environments. Ecology and Evolution, 12(3), e8684. 10.1002/ece3.8684

