## supplementary material for "Multiple within species comparisons show Tanganyikan cichlid fish have larger brains in less structurally complex habitats"

**Curvature Computation and Analysis**

In the context of habitat complexity assessment, the curvature of the habitat surface plays a crucial role in quantifying structural variation. Curvature, a geometric property of a surface, describes how much a surface bends at a given point (**Figure S1**). For this study, we computed curvature information based on the discrete triangular surfaces generated from our 3D reconstructions (see main text). Curvature was analyzed to capture local surface complexity, which is not easily detectable using simpler metrics such as rugosity.

The curvature information was derived using the **maximum principal curvature** of each triangle on the surface. Two key metrics were considered: **Max Curvature** and its inverse, **Inverse Curvature**. The decision to use Inverse Curvature was based on several comparative analyses of different parameters and their effectiveness in capturing the detailed surface geometry.

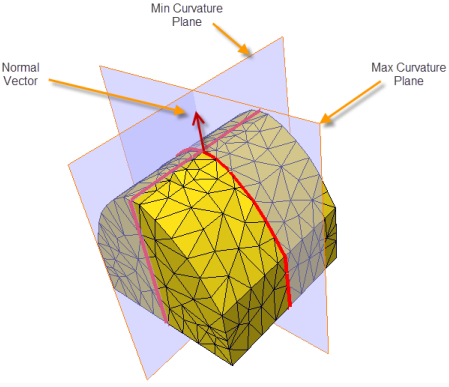

**Figure S1.** Schematic representation of curvature calculation. The Max Curvature at a point on the surface describes the sharpest bending direction, while the Normal Vector represents the perpendicular direction to the surface. The Max and Min Curvature planes intersect the surface at different angles, and the Inverse Curvature, used in this study, represents the radius of the sphere that locally fits the surface.

**Why Inverse Curvature Was Chosen**

Inverse Curvature was selected as a key metric for the following reasons:

*Detailed Local Surface Information*

Unlike rugosity, which provides a global measure of surface roughness, Inverse Curvature allows us to capture local surface details at a finer scale. It provides a point-by-point assessment of how the surface curves in different regions, offering insight into small-scale structural features that may not be visible through rugosity alone.

*Surface Uniformity and Complexity*

By calculating Inverse Curvature, we can quantify the degree of surface uniformity. Larger values of Inverse Curvature indicate flatter areas, while smaller values represent areas with sharp features, such as rocks or steep inclines. This allows us to assess both overall smoothness and localized complexity within each habitat. For example, habitats with a higher concentration of sharp surface features will have a broader distribution of Inverse Curvature values, indicating greater structural complexity.

*Physical Interpretation of Surface Geometry*

Inverse Curvature has a direct physical interpretation in terms of the radius of a locally fitting sphere. This provides a meaningful way to compare surface features across different habitats, as it relates directly to the geometry of the surface. In practical terms, the radius can be thought of as how "curved" or "flat" a surface feature is—habitats with smaller radii (i.e., smaller Inverse Curvature values) will have more pronounced bends, while habitats with larger radii (larger Inverse Curvature values) will be flatter.

*Complementing Rugosity and Height Distribution*

Inverse Curvature complements rugosity and height distribution by adding an additional layer of detail regarding surface structure. While rugosity captures the overall roughness of the habitat and height distribution reflects topographical variation, Inverse Curvature allows for a more localized analysis of surface bending and curvature. This provides a more comprehensive picture of habitat complexity, as different habitats may have similar rugosity values but vary significantly in terms of local surface features and curvature.

*Quantifying Landmark Distribution*

Finally, Inverse Curvature helps in assessing the distribution of surface landmarks such as rocks and other prominent features. By examining the probability distribution of Inverse Curvature values within each habitat, we can determine the spatial uniformity and the presence of localized surface irregularities. This allows us to identify habitats with a more homogeneous distribution of surface features versus those with more dispersed or concentrated landmarks.

**Habitat Complexity as the Key Factor**

Through extensive modelling in this study, while there are numerous confounding factors that could contribute to brain volume variation, we identified habitat complexity as the predominant factor influencing brain size and architecture.

Firstly, brains are energetically costly organs (Niven et al., 2007; Navarrete et al., 2011; Kotrschal et al., 2013). It is conceivable that fish with limited food intake might have smaller brains. However, we considered and rejected the hypothesis that the larger brain size observed in fish from less complex habitats (such as Habitat C) might result from greater energy availability. This is because body condition (mass adjusted for standard length) did not differ significantly between fish from these habitats. Additionally, research has shown that more complex habitats are generally associated with better foraging opportunities. Therefore, it seems unlikely that the larger brain sizes in the less complex Habitat C habitat are a result of higher energy availability. Secondly, other studies have demonstrated that factors such as water temperature (Gillooly & McCoy, 2014; Yu et al., 2014; Závorka et al., 2020), visibility, and water depth can influence brain size. However, the habitats we selected for this study are geographically very close to each other, and during the fieldwork, water temperature and visibility were consistent across these locations, despite daily fluctuations. While water depth varied among the habitats, our original linear mixed model (LMM) included water depth as a factor, and it was found to be non-significant. Moreover, Habitat C, the habitat associated with the largest brains, was neither the deepest nor the shallowest, further indicating that water depth is unlikely to be a major contributor to brain size variation. Thirdly, the random selection of species and sex in our study could have influenced brain size patterns if species distribution or sex ratios differed significantly across habitats. However, through a series of model selections, we confirmed that sex is not a significant predictor. Finally, habitat complexity emerged as the significant predictor in our initial model. Moreover, when we classified habitats based on their complexity—labelling Habitat C as "low complexity" and Habitat A, Habitat B, and Habitat D as "high complexity"—the inclusion of complexity as a predictor significantly improved model performance. This analysis strongly suggests that differences in habitat complexity drive the observed variation in brain size.

**Sex and water depth did not influence brain volume**

We investigated whether the observed variability in brain volume among habitats was attributable to habitat complexity or other confounding factors. We confirmed that habitat complexity was the predominant predictor. Models incorporating sex (F = 0.0759, p = 0.7833) and water depth (F = 0.0683, p = 0.7942) as predictors did not significantly affect brain volume nor improve model performance. Following reclassification of habitats into a new factor labelled "habitat complexity," where Habitat C was designated as "low" complexity and Habitat A, Habitat B, and Habitat D as "high" complexity, we noted improved model performance. This was evident from significant reductions in the Akaike Information Criterion (ΔAIC = 8.74075) and the Bayesian Information Criterion (ΔBIC = 14.638271), validating the inclusion of habitat complexity in further analyses. Additionally, Model 6, which analysed body weight as a function of complexity, standard length, and random effects by species, showed no significant differences in body weight adjusted for standard length between habitats of high and low complexity (F = 0.1262, p = 0.7229). This outcome indicates that fish across varied habitat complexities employ similar energy allocation strategies, as illustrated in **Figure S2**.

**Table S1. Summary of samples used in this study**

|  | Habitat site | | | | |
| --- | --- | --- | --- | --- | --- |
| Species | Habitat B | Habitat A | Habitat C | Habitat D | Total |
| *Altolamprologus compressiceps* | 1 | 1 | 3 | 3 | 8 |
| *Eretmodus cyanosticus* | 0 | 4 | 5 | 6 | 15 |
| *Interochromis lookii* | 4 | 0 | 3 | 2 | 9 |
| *Julidochromis ornatus* | 5 | 5 | 5 | 5 | 20 |
| *Lepidolamprologus elongatus* | 4 | 0 | 2 | 0 | 6 |
| *Neolamprologus pulcher* | 6 | 3 | 5 | 7 | 21 |
| *Neolamprologus tetracanthus* | 5 | 4 | 6 | 5 | 20 |
| *Telmatochromis temporalis* | 4 | 6 | 6 | 5 | 21 |
| *Tropheus moorii* | 1 | 2 | 3 | 3 | 9 |
| *Variabillichromis moorii* | 4 | 3 | 1 | 4 | 12 |
| Total | 34 | 28 | 39 | 40 | 141 |

**Table S2. One time survey of ecological features of four selected habitats**

| Habitat | Habitat C | Habitat D | Habitat B | Habitat A |
| --- | --- | --- | --- | --- |
| Predators | 2 | 1 | 7 | 0 |
| Competitors | 57 | 81 | 77 | 42 |
| Water Depth | 8.7m | 7.6m | 11.5m | 5.3m |

**Table S3: Effects of environmental complexity on brain volume across species**

| **Species** | **Complexity Estimate** | **Std. Error** | **t-value** | **p-value** |
| --- | --- | --- | --- | --- |
| *Altolamprologus compressiceps* | 0.06708 | 0.10450 | 0.642 | 0.5492 |
| *Eretmodus cyanosticus* | 0.11360 | 0.10659 | 1.066 | 0.307485 |
| *Interochromis lookii* | 0.11927 | 0.12405 | 0.961 | 0.37345 |
| ***Julidochromis ornatus*** | 0.24873 | 0.09451 | 2.632 | **0.0175** |
| *Lepidolamprologus elongatus* | 0.1747 | 0.3478 | 0.502 | 0.6500 |
| *Neolamprologus pulcher* | 0.12482 | 0.06631 | 1.882 | 0.076 |
| ***Neolamprologus tetracanthus*** | 0.14692 | 0.05898 | 2.491 | **0.0234** |
| ***Telmatochromis temporalis*** | 0.28787 | 0.10841 | 2.655 | **0.0161** |
| ***Tropheus moorii*** | 0.5779 | 0.1575 | 3.669 | **0.010468** |
| *Variabillichromis moorii* | 0.21118 | 0.21903 | 0.964 | 0.360164 |

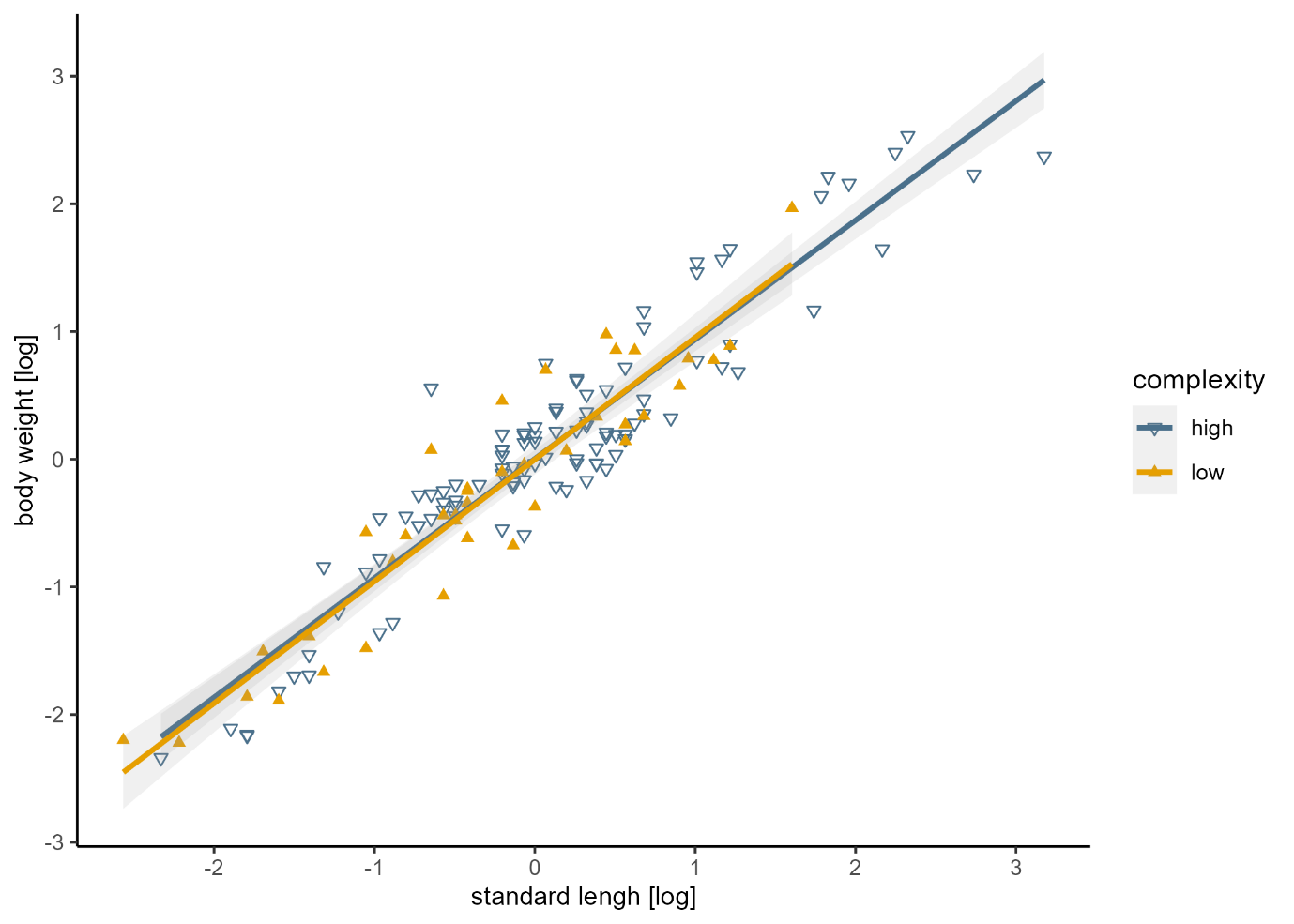

**Figure S2.** The relationship between adjusted body weight and standard length (both log-transformed), along with the linear fits for each habitat. There is no significant difference between these two linear fits (t = -0.355, p = 0.723).
